# δ-Conotoxin Structure Prediction and Analysis through Large-scale Comparative and Deep Learning Modeling Approaches

**DOI:** 10.1101/2024.05.30.596722

**Authors:** Stephen McCarthy, Shane Gonen

## Abstract

The δ-conotoxins, a class of peptide produced in the venom of cone snails, are of interest due to their ability to inhibit the inactivation of voltage-gated sodium channels causing paralysis and other neurological responses, but difficulties in their isolation and synthesis have made structural characterization challenging. Taking advantage of recent breakthroughs in computational algorithms for structure prediction that have made modeling especially useful when experimental data is sparse, this work uses both the deep-learning based algorithm AlphaFold and comparative modeling method RosettaCM to model and analyze 18 previously uncharacterized δ-conotoxins derived from piscivorous, vermivorous, and molluscivorous cone snails. The models provide useful insights into structural aspects of these peptides and suggest features likely to be significant in influencing their binding and different pharmacological activities against their targets, with implications for drug development. Additionally, the described protocol provides a roadmap for the modeling of similar disulfide-rich peptides by these complementary methods.

## 1. Introduction

Conopeptides are peptides that have been evolved by marine cone snails to protect against predators and to capture prey. Each species of cone snail may produce over 1,000 different peptides^[1]^ and estimates for the total number of conopeptides range from 50,000 to millions,^[2,3]^ but structural characterization remains sparse. The targets for these conopeptides are varied and include several protein classes of significance in human disease such as ion channels and G protein-coupled receptors (GPCRs). Although conopeptides rarely comprise more than 40 amino acids, individual conopeptides can show high levels of target specificity and potency, including, in some cases, the ability to distinguish between receptor subtypes.^[4]^ These properties have made them the subject of considerable research, both as tools in neurological research and as leads for drug development.

Conotoxins are conopeptides that exert a toxic effect on the envenomed organism and are classified according to their target and pharmacological effects. δ-conotoxins (**Figure 1A**) interact with voltage-gated sodium channels (Na_v_s), where they inhibit the fast inactivation phase of channel gating and prolong an open channel conformation, similar to scorpion α-toxins.^[5–7]^ While the target subtype selectivity of most δ-conotoxins remains undetermined, intriguing differences have been noted between δ-conotoxins derived from molluscivorous (mollusk-eating) cone snails compared with those from piscivorous (fish-eating) cone snails (**Figure 1B**): while δ-conotoxins from piscivorous cone snails show activity against vertebrate Na_v_s, peptides from molluscivorous snails (with the exception of Am2766^[8]^) do not show activity on vertebrate neurons, while retaining their characteristic activity against mollusk Na_v_s.^[4,5,9,10]^ Radioligand binding studies on the molluscivorous cone snail peptide TxVIA nonetheless suggest that it retains the ability to bind to vertebrate Na_v_s, but without having any pharmacological effect on the channel.^[5]^ A structural basis for this ‘silent binding’ has not yet been determined, and experimental structures of δ-conotoxins to date are limited to two peptides from molluscivorous cone snails, TxVIA and Am2766, and one from a piscivorous cone snail, EVIA, which has been the subject of several structural studies.^[11–14]^ The peptide EVIA is an outlier among conopeptides from piscivorous snails due to its longer loop 2 (see **Figure 1A** for nomenclature), and consequently little structural information exists for most members of this class.

**Figure 1.**
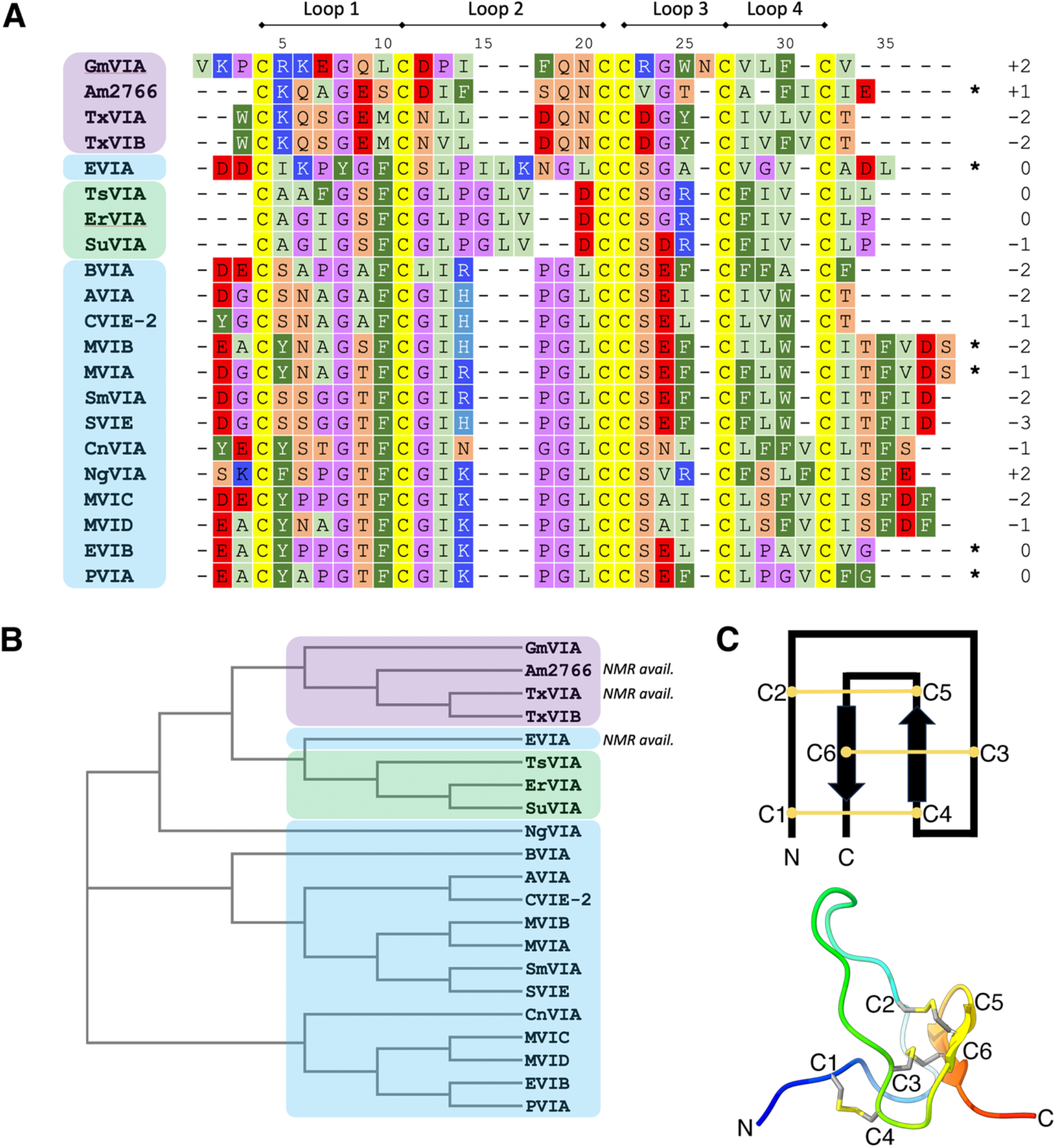
A) Sequence alignment of the δ-conotoxins modeled in this study, together with the overall net charge and loop nomenclature. Peptides from piscivorous, vermivorous, and molluscivorous cone snails are highlighted in blue, green, and purple, respectively. C-terminal amides are indicated with *. B) Phylogenetic analysis of the sequences shown in A indicating peptides with available experimental structures and colored according to the color scheme in A. C) (top) Features typical of an inhibitor cystine knot (ICK) peptide, showing the connectivity of the disulfide bonds and (bottom) One conformation of the NMR-derived structure of δ-EVIA (PDB ID: 1G1P)^[13]^, showing the classic ICK fold.

Here we adopt a computational modeling approach to determine three-dimensional models of all δ-conotoxins without an experimental structure, using the complementary modeling tools AlphaFold and RosettaCM. AlphaFold is a deep learning based algorithm that generates models using multiple sequence alignments (MSAs) and structural data as inputs,^[15]^ whereas RosettaCM is a threading modeling method requiring one or more template structures.^[16]^ Computational modeling is especially useful for studies of δ-conotoxins, as the complex inhibitor cystine knot (ICK) disulfide bonding pattern (**Figure 1C**), low oxidative folding yields and high hydrophobicity have hampered the synthetic efforts necessary for structural studies. Analyzing multiple models generated by both AlphaFold and RosettaCM provides insights into key structural features of the δ-conotoxins and the bases for their pharmacological properties, assists in developing modeling strategies for disulfide-rich peptides, and aids in the characterization of δ-conotoxins yet to be discovered.

## 2. Results

Using AlphaFold, 5 decoys per target δ-conotoxin were generated, and all 5 models were included in the analysis. For the RosettaCM models the lowest-energy representatives of the top 5 clusters, generated from 5,000 total models, were used for analysis. Representatives for each peptide, together with corresponding quality metrics, are highlighted in **Figure 2**, and our analysis and observations from these models are described below.

**Figure 2.**
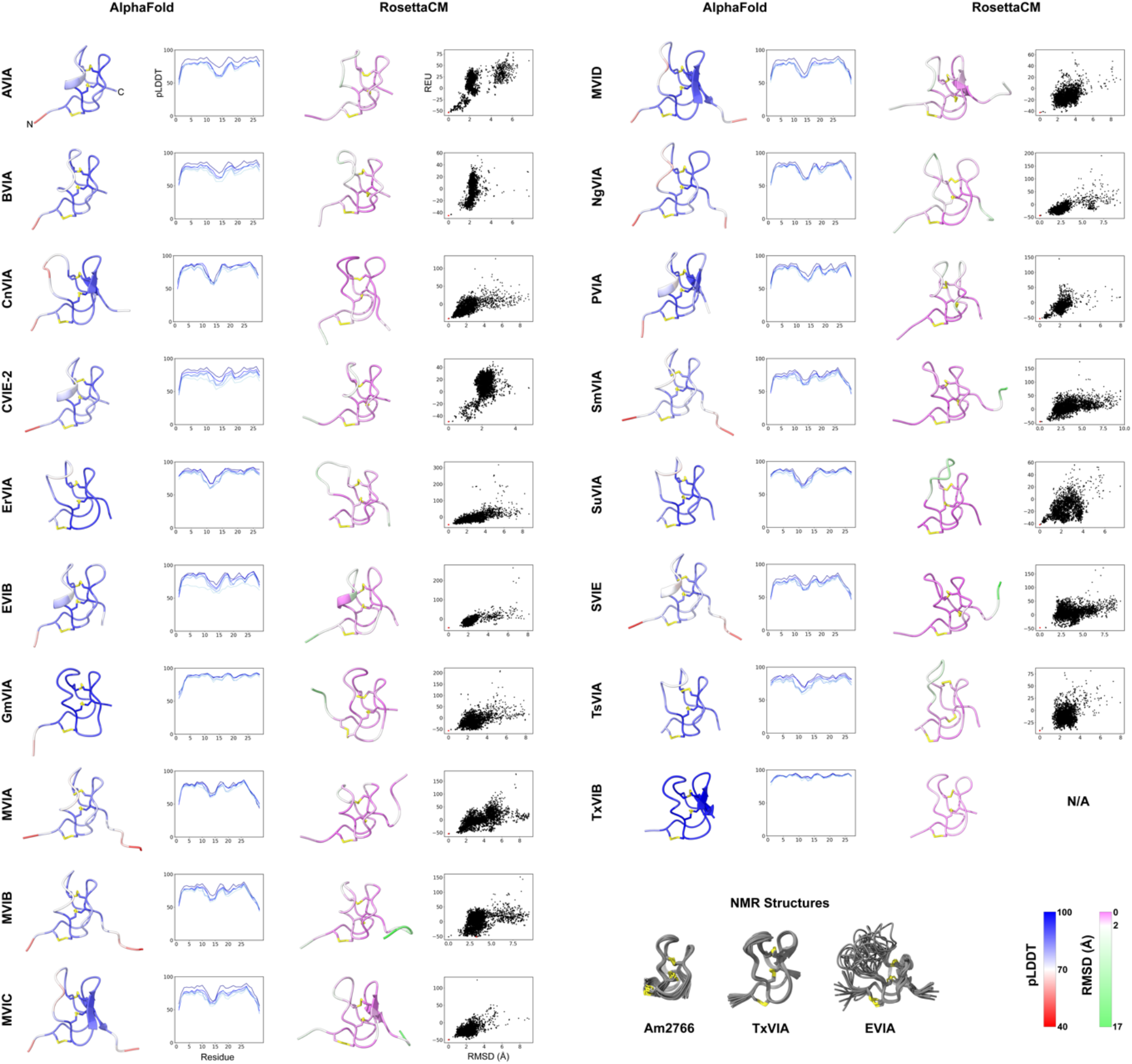
Results of the modeling pipeline for all peptides. Disulfide bonds are highlighted in yellow. From left: best-scoring model produced by AlphaFold, colored by per-residue pLDDT score; plot of pLDDT per residue for the models generated by AlphaFold. Models with higher overall pLDDT scores are in darker blue; best-scoring model produced by RosettaCM, colored by per-residue root-mean-square deviation (RMSD) to representatives of the top 5 best-scoring clusters; score-RMSD funnel plot for all 5,000 decoys produced by RosettaCM, with score measured in Rosetta energy units (REU) and RMSD calculated by distance to the lowest-energy model, and with the displayed model indicated in red;. Below: experimentally-determined NMR ensembles used as templates in this study. RMSD and pLDDT color bars are shown bottom right.

### 2.1. Peptides show the expected ICK geometry

Peptide models were initially evaluated by checking to see that they had formed the expected ICK fold with the correct disulfide connectivity (C1-C4, C2-C5, C3-C6) (Figure 1C); in the case of the RosettaCM models this geometry was enforced and so all models met this criterion. For AlphaFold models, however, no disulfide bonding pattern was stipulated in the inputs; nonetheless AlphaFold produced highly accurate predictions, with only a single model (out of a total of 90) displaying incorrect disulfide connectivity (**Figure S1**). Notably, an attempt to perform modeling in AlphaFold using only MSA data and without the addition of template structures as inputs resulted in considerably poorer results, with approximately half the target peptides showing no models with the correct ICK topology (data not shown).

### 2.2. Peptide termini are generally flexible

For our purposes, the N- and C-terminal regions are defined as the residues prior to the first cysteine residue, and after the last cysteine residue, respectively. Consistent with experimental evidence showing that these regions are largely unstructured in ICK peptides,^[13]^ we observe decreased pLDDT scores in the AlphaFold models, which is ascribed to regions that are either intrinsically disordered or ordered only in complex (**Figure S2**).^[15,17,18]^ We also observe a high degree of divergence in the RosettaCM models of the termini, and correspondingly higher per-residue root-mean-square deviation (RMSD) scores.

### 2.3. Secondary structure is limited

The ICK fold classically contains a short β-turn near the C-terminus that incorporates C5 and C6; very occasionally a short third strand is observed around C2.^[19]^ Our models largely conform to this pattern, with limited secondary structure aside from this C-terminal β-turn. However, one RosettaCM model contains a formal helical half-turn just prior to C3, and this turn is also present in 5/18 of the top-ranked AlphaFold models, all of which are derived from piscivorous cone snails (Figure 2). This element is unusual in experimental structures of ICK peptides; an example featuring a helix in this position is the insecticidal funnel-web spider toxin U21-hexatoxin-Hi1a, however this peptide contains a number of additional structural features beyond the ICK scaffold core and the helix in this section of the peptide might be more properly considered to be an insertion into the otherwise unstructured loop.^[20]^ Since this region of the δ-conotoxins is relatively flexible (see below) it is possible that the observed secondary structure in this region is transitory and does not represent a significant population in solution, but it is also possible that conotoxins with a longer loop 2 permit formation of short secondary structure elements that would not otherwise be observed.

### 2.4. Peptide flexibility is greatest in loop 2

Of the four disulfide-constrained loops, loop 2 consistently shows the lowest mean pLDDT score for the AlphaFold models and the highest RMSD between RosettaCM models (Figure S2), strongly suggesting that this region is flexible. This flexibility can be observed in the ensemble of structures for EVIA determined by NMR (Figure 2); while this peptide was used as a template in our RosettaCM modeling, only a single conformation was used and consequently all the structural diversity observed in this loop has been independently derived by the modeling algorithms.

While loop 2 is by far the longest loop in EVIA (9 amino acids), most of the other conotoxins in this study have a shorter loop 2 (6 amino acids) with the exceptions being ErVIA, SuVIA, and TsVIA from vermivorous snails (7 amino acids). It is therefore notable that the other peptides retain flexibility in this loop despite a shorter loop length; loop 1 has considerably lower predicted RMSDs than loop 2 (Figure S2), despite also consisting of 6 amino acids in all cases.

This region is comparatively rigid in δ-conotoxins from molluscivorous snails. Examining the predicted structures for GmVIA reveals a role for the sidechain of Q16, which can form hydrogen bonds to the backbone carbonyl of P13 and the backbone amide of C24 (**Figure S3A**). The AlphaFold models additionally show hydrogen bonding to the C24 backbone carbonyl. This glutamine is conserved across all δ-conotoxins from molluscivorous peptides, and similar hydrogen bonding arrangements are found in some conformations of Am2766 (**Figure S3B**) and TxVIA.^[11,12]^ These peptides also lack the glycine immediately following C2 that is conserved across most δ-conotoxins from piscivorous cone snails and which could contribute to the additional flexibility of loop 2 in these peptides.

### 2.5. Not all peptides show a continuum of positions in loop 2

In some models we do not observe a continuum of positions for loop 2 across the full range of motion, as might be expected for a disordered region where there are low energetic barriers to movement. In these cases, which are restricted to some δ-conotoxins from piscivorous cone snails, we observe that the top-ranked models are clustered around a few discrete positions (**Figure S4**). This suggests that the loop may be able to interconvert between different populations, with implications for bioactivity (see Discussion).

### 2.6. A structural role for hydroxyproline residues in RosettaCM models

Post-translational modifications are common among conotoxins from all functional classes. Among the δ-conotoxins in this study, 4-hydroxyproline is modeled at position 6 (in the middle of loop 1) in five peptides, and at position 14 (middle of loop 2) in 12 peptides. Of these two Hyp residues, the first has a clear structural rationale: the additional hydroxy group on the sidechain of Hyp6 forms a hydrogen bond with the sidechain of the conserved S19, where it mitigates any flexibility of loop 1 as well as further stabilizing S19, which has a role in forming the tight turn at the start of loop 3 (**Figure S3C**).

While this Hyp residue has a clear structural justification, the same cannot be said for the Hyp residue on loop 2. This loop is flexible (see above) but in all conformations the hydroxyl group is directed away from the rest of the peptide where it cannot form direct intramolecular interactions. This raises the possibility that it either contacts the target Na_v_ or influences the structure less directly.

### 2.7. δ-Conotoxins with an extended loop 2 show a mixture of cis- and trans-proline

Cis-prolines are uncommon in protein structures due to the increased energy of this conformation relative to trans-proline, and their prevalence is estimated to be approximately 6% of all prolines.^[21]^ It is therefore significant that in the RosettaCM models of two δ-conotoxins in this study with an extended loop 2, a conserved Pro at position 11 shows a mixture of cis and trans conformations in the 5 best-scoring clusters (**Figure S5**). This result extends published NMR studies of EVIA, which demonstrated a mixture of cis- and trans-prolines at the equivalent position in an approximately 1:1 ratio with slow interconversion.^[13,14]^ One NMR study also demonstrated that a P13A EVIA mutant produced a 100% trans conformation at this bond, and also halved the potency of the peptide compared to the wild type in a competitive binding assay using ^125^I-labeled TxVIA on rat brain synaptosomes.^[13]^ This suggests that either the sidechain makes direct contact with the target, or that the cis-proline conformation is important for positioning other interacting residues.

Notably, the cis-proline bonds are only found in the RosettaCM models; the equivalent AlphaFold models show only the trans conformation. Comparing the top-scoring cluster for TsVIA (which has a cis-proline at position 11) with the corresponding top-ranked AlphaFold model shows that the cis-proline bond allows the L10 and V14 sidechains to project towards loop 3 and be partially buried in the peptide core (**Figure S3D**). By contrast, the trans-proline conformation projects the sidechains away from the peptide core where they are fully exposed to solvent; it can therefore be hypothesized that the cis-trans isomerization at proline in this loop balances the higher-energy cis-conformation with the energetic penalty from fully-exposed hydrophobic leucine and valine sidechains.

### 2.8. A consensus surface-exposed hydrophobic patch

The presentation of hydrophobic sidechains on the solvent-accessible surface is disfavored in most protein structures but is a common feature of ICK peptides; it has been hypothesized that the reason for forming such a complex, covalently-linked peptide core is to force exposure of these residues, even against the energetic penalty.^[22–24]^ These hydrophobic patches are often important for binding.^[25,26]^ Examining the available experimental structures of δ-conotoxins, together with our models, we find a common hydrophobic patch on the peptide surface (**Figure S6**). Sequence alignments of the δ-conotoxins show that these regions are uniformly hydrophobic, across all types of cone snail. This consensus strongly suggests that these residues are important for binding of the peptides to the target, including the ‘silent binding’ by the δ-conotoxins from molluscivorous cone snails.

### 2.9. Some RosettaCM models of piscivorous cone snail δ-conotoxins show a cis-nonPro bond in loop 2

Several of the top-ranked RosettaCM models (but none of the equivalent AlphaFold models) contained a non-proline cis-peptide bond at the conserved glycine-isoleucine motif in loop 2 immediately following the C2-C5 disulfide bond. Peptides that lacked the GI motif, such as GmVIA and TxVIB (derived from molluscivorous cone snails) and BVIA (from a piscivorous snail), did not contain a cis-peptide; neither did any of the peptides derived from vermivorous snails (TsVIA, ErVIA and SuVIA) even though these peptides contain a similar GL motif as part of a longer loop 2. Given the rarity of this structural feature in experimentally derived protein structures, we repeated the RosettaCM modeling protocol with a further two different score functions to investigate methodological explanations for its appearance in our model set (see Methods), only to obtain similar results (**Table S1**). Although none of the AlphaFold models contained non-proline cis-peptide bonds, in some instances a twisted (non-planar) peptide bond was observed in the C-terminal regions which suggests some difficulties in modeling correct geometries in regions of higher uncertainty (see Discussion).^[27]^

## 3. Discussion

In this study we calculated models for δ-conotoxins without experimental structures, using both AlphaFold and RosettaCM. The models show many structural features that are well-known properties of all δ-conotoxins, such as the disulfide bond connectivity typical of ICK peptides and surface-exposed hydrophobic patch that arises from the rigidity of this network. However, several differences are observed between the δ-conotoxins derived from piscivorous cone snails compared with those from molluscivorous cone snails, which provide suggestions for the structural basis of the ‘silent binding’ phenomenon.

The most notable difference between the two classes of peptides is the flexibility observed for loop 2, which is seen in all δ-conotoxins from piscivorous and vermivorous cone snails, but not among the δ-conotoxins from molluscivorous snails (Figure S2). A previous structure-activity relationship study of PVIA noted that the mutants P9A and I12A – both highly conserved residues in δ-conotoxins from piscivorous snails but not molluscivorous snails – retained the ability to bind to Na_v_s but lost their pharmacological properties.^[9]^ Since these residues are located on loop 2, the flexibility of this loop observed in our models raises the question of how these residues are positioned to interact with the target. Since F9 is located on loop 1, the AlphaFold models show a largely consistent position for the sidechain. I12 shows greater variance; in some models it is orientated away from loop 2 and the disulfide-bonded core and closer to the F9 sidechain, whereas in others it is directed inwards and positioned closer to loop 3. We speculate that, given the relative exposure to solvent of the I12 sidechain in the vertical position, as well as its closer positioning to the bioactive residues F9, the former orientation is more likely to be bioactive.

Examining the sequences of other classes of peptide known to elicit similar pharmacological properties when binding to Na_v_s is also illustrative. A study of Australian funnel-web spiders uncovered 22 peptides of the δ-hexatoxin class.^[28]^ These peptides similarly inhibit the inactivation of voltage-gated sodium channels (including in vertebrates) and are thought to share a similar binding site to the δ-conotoxins; although they share a common ICK disulfide bonding core, the δ-hexatoxins are typically larger and possess an additional disulfide bond. Sequence alignments of 14 representative δ-hexatoxins with the δ-conotoxins in this study show that an aromatic residue is conserved at the equivalent of F9 in all peptides (typically it is tryptophan), apart from the δ-conotoxins from molluscivorous cone snails, strongly implicating this residue in bioactivity (**Figure S7**).

The δ-conotoxins have a low net charge, ranging from -3 to +2 (Figure 1A); this contrasts with other peptide venom toxins, such as the μ-conotoxins or spider venom ICK peptides, which often have a very high positive net charge.^[29]^ This likely contributes to the low aqueous solubility and refolding yields of these peptides, which has hindered their structural and functional characterization. Since a high net positive charge has been shown to be an important determinant of potency against voltage-gated sodium channels for related tarantula-derived ICK peptides due to a membrane-first binding mechanism,^[30,31]^ the low charge of the δ-conotoxins would suggest that a different binding mechanism applies.

Although both piscivorous and molluscivorous snail-derived δ-conotoxins share the low overall charge, the charged residues are distributed differently across the loops, and recent docking studies have implicated these charged residues as determinants of channel binding.^[14,32]^Piscivorous snail δ-conotoxins usually have acidic residues at the N-terminus, on loop 3, and sometimes at the C-terminus, while the basic residues are limited to a single position on loop 2 (Figure 1a). Molluscivorous snail-derived δ-conotoxins, however, universally have a basic residue immediately after C1, while acidic residues are located near the end of loop 1, and immediately after C2. In cases where these residues are not negatively charged a polar residue is often found in its place. The conserved positioning of the basic residue in peptides from piscivorous snails, in a loop that is flexible and known to contain residues important for the inhibitory abilities of the peptide, suggests a role in the mechanism of action of these peptides. The supposed binding site for δ-conotoxins^[6]^ (above voltage-sensing domain (VSD) IV of voltage-gated sodium channels) permits access to the ladder of charged amino acids within VSD IV that are responsible for the movement of the domain on activation. Disruption of this ladder by basic residues is a known mechanism of channel inhibition by other venom peptides, and could play a role in inhibiting inactivation.^[33–35]^

Related to the positioning of these residues on loop 2 is the cis/trans-isomerization at the conserved proline on this loop among δ-conotoxins from piscivorous snails. The appearance of the cis-isomer is restricted to RosettaCM models of peptides with an extended loop 2 (i.e. longer than 6 residues) (Figure S5), suggesting that the cis-conformation cannot form in the tighter and more constrained 6-residue loop that is most prevalent for loop 2. One possible functional explanation for the formation of the cis-isomer was suggested by Tietze et al.,^[14]^ who hypothesized that the cis conformation was likely to increase binding affinity by presenting both a hydrophobic side chain and a backbone carbonyl (capable of accepting a hydrogen bond) to the Na_v_.

All AlphaFold models showed exclusively trans-proline. To our knowledge a systematic study of the propensity of AlphaFold to predict cis-prolines has not yet been performed, but in CASP14^[36]^ it correctly predicted 42 trans-prolines and 3 out of 4 cis-prolines,^[37]^ suggesting that AlphaFold performs well at identifying cis-prolines in its targets. One explanation for the results seen in our models is that the cis-proline peptide bond can interconvert between conformations, as has been experimentally observed for EVIA, and the trans conformation is selected for statistical reasons.

Cis-peptide bonds at residues other than proline are extremely unusual in protein structures and their appearance in our set of best-ranked RosettaCM models prompted further scrutiny. To verify the initial models we repeated the modeling using two further scorefunctions and an alternative refinement strategy, but nonetheless obtained similar results across both new datasets. The non-proline cis-peptides seen in the top-ranked models all occur at the same position – the glycine-isoleucine bond immediately following the C2-C5 disulfide bond in conotoxins derived from piscivorous cone snails. Peptides that lack this motif or that have a longer loop 2 do not have cis-peptide bonds anywhere in the model. These results are consistent with the observation that approximately 80% of characterized cis-Xaa-nonPro bonds have glycine prior to the cis-peptide bond,^[38]^ since the steric penalty is minimized when the clashing sidechain is a proton. Despite their rarity, non-proline cis-peptide bonds have been characterized in NMR structures of α-scorpion toxins;^[39–41]^ these peptides similarly inhibit Na_v_ inactivation and also bind to the extracellular loops of VSD IV.

As previously noted, the AlphaFold models do not contain any cis-peptide bonds. However, in a handful of cases the AlphaFold models have ‘twisted’ peptide bonds (ω > ± 30° deviation from planar) which are practically nonexistent in experimental protein structures (Table S1). It has been shown that AlphaFold is more likely to predict these model geometries in low-confidence regions (pLDDT < 60)^[27]^ and this is also observed in our dataset, with the twisted peptides exclusively occurring in the disordered C-termini; similar features are seen in models of these peptides in the AlphaFold database.^[17]^ Nonetheless the conclusions in this paper are solely derived from models without cis or non-planar peptide bonds.

In this work we use both a deep learning algorithm (AlphaFold) and a conventional homology and energy minimization strategy (RosettaCM) to generate our model sets. Each method has its strengths and weaknesses when modeling these complex peptides. Threading algorithms are at their most accurate when modeling sequences with high homology to the available templates, as is often the case with ICK peptides given the universality of this framework across different species; RosettaCM is particularly well suited for this given its ability to use multiple template structures simultaneously. Conotoxins often have extensive post-translational modifications; C-terminal amides and 4-hydroxyproline residues were present in our target set but other conotoxin classes are known to have different modifications, including D-amino acids, γ-carboxylation of glutamic acid residues, and L-6-bromination of tryptophan residues.^[42]^ Rosetta has an extensive library of patches that enable these modifications to be handled natively during modeling which offers a significant advantage.

AlphaFold is a newer modeling algorithm that uses deep learning methods to calculate the structure using an MSA, residue-pair interaction data, and, optionally, experimental structures as templates. AlphaFold’s chief advantage is its accuracy: in the CASP assessments of protein structure modeling methods in which AlphaFold has participated it considerably outperformed other algorithms.^[36]^ It also has the advantage that it can predict structures based on the sequence data alone, given a sufficiently large MSA, and does not entirely rely on experimental structures as homology methods do. Given the challenges in experimental characterization presented by certain conotoxins, this could significantly improve model quality. While the original AlphaFold algorithm has been extended to improve modeling of protein-protein and protein-ligand complexes,^[43]^ a full treatment of post-translational modifications is not available; this presents a drawback in the modeling of conotoxins given the frequency and importance of post-translational modifications in conotoxin structure and function.

## 4. Conclusion

Our computational study has expanded the range of δ-conotoxins with modeled structures, including, for the first time, a class of δ-conotoxins from piscivorous cone snails for which experimental structures are sparse and challenging to obtain. We note intriguing similarities and differences between peptides originating from piscivorous, molluscivorous, and vermivorous cone snails and identify features likely to be of relevance to their binding and activity against their target Na_v_s, including the ‘silent binding’ phenomenon. Our protocol additionally suggests a roadmap for modeling similar disulfide-rich peptides. Further structural characterization of these δ-conotoxins, both in isolation and in complex, will shed additional light on their mechanism of action and provide new avenues for rational drug design.

## 5. Methods

### Database searching, template selection and preparation

δ-conotoxins were identified using ConoServer.^[44,45]^ Peptides annotated with the cysteine framework VI/VII and pharmacological family δ were selected; synthetic constructs were excluded. Three peptides, Am2766, TxVIA, and EVIA, have experimental structures and were designated as templates for modeling the remaining δ-conotoxins in RosettaCM.^[11–14]^ 4-Hydroxyproline residues were designated as prolines for sequence alignments. Due to the very high sequence similarity to TxVIA, conotoxin TxVIB was modeled using a different protocol (see below).

All template peptide structures were determined by nuclear magnetic resonance (NMR) and therefore comprise an ensemble of structures consistent with the calculated restraints. To prepare a single input model per template for RosettaCM, the ensemble structures were separated into individual PDB files and cleaned using the clean_PDB.py script supplied with Rosetta. For 1YZ2, which has a C-terminal amide, the OXT atom was replaced in one round of Rosetta FastRelax, then edited to restore a C-terminal amide. All structures were then energy-minimized (5 models per input structure) using Rosetta Relax,^[46,47]^ and the lowest-energy model was selected as the template.

### Modeling with AlphaFold

AlphaFold was installed from source (2021-07-14 parameters) and modeling was performed with the –max_template_date=2021-09-01 and –preset=casp14 flags, and the use of experimental structures as templates was permitted. Hydroxyproline residues were modeled as unmodified prolines. The five ranked models produced were assessed for model quality by their per-residue pLDDT scores and model geometry was assessed with MolProbity.^[48]^

### Model generation in RosettaCM

Sequences were obtained from ConoServer. Since sequence data for some of these peptides is obtained through genomic analyses, which do not indicate the presence or absence of post-translational modifications, hydroxyproline residues and C-terminal amides were modeled where annotations in ConoServer indicated their likely presence; note that some of these may be derived by similarity and do not guarantee a modification at that position.

Sequence alignments were generated using ClustalΩ^[49]^ and adjusted where necessary to ensure alignment of the cysteine residues. The phylogenetic tree (Figure 1B) was calculated using the phylogeny tool ClustalW2.^[50,51]^ Initial threaded models were calculated for each peptide using the setup_RosettaCM.py script provided with Rosetta,^[16]^ then 1,000 initial models were calculated using the rosetta_cm.xml script. This script was modified where necessary to incorporate 4-hydroxyproline residues and C-terminal amides using the ModifyVariantType mover. The energy of the models was minimized using the Relax application with the current ref2015 scorefunction,^[52]^ generating 5 relaxed models per input model for a total of 5,000 models per peptide. The -fix_disulfs flag was used, together with a parameters file defining the disulfide bond connectivity, to ensure formation of the correct cysteine framework.

This protocol was similarly followed when modeling using the cartesian scorefunctions beta_nov16_cart and ref2015_cart,^[53]^ except the number of output hydridized models was set to 5,000 and the -relax:dualspace flag was included to instigate dualspace refinement.

All scripts and flags used are provided as Supplementary Material.

### Model clustering and scoring

Models were clustered using Rosetta’s energy-based clustering algorithm,^[54]^ with a cluster radius of 1 Å. The 5 lowest-energy clusters were used for analysis, and decoys were aligned to the lowest-scoring model in ChimeraX.^[55,56]^

### Modeling of TxVIB

The sequence of TxVIB is identical to that of TxVIA except for two mutations (L11V/L24F).^[57,58]^ Modeling of TxVIB was therefore carried out using the mutagenesis tool in PyMOL^[59]^ to mutate the two residues, followed by energy minimization using Rosetta Relax (500 models). The lowest-scoring model was selected for analysis. The standard AlphaFold modeling protocol was used for TxVIB.

Figures were produced in ChimeraX, PyMOL, Microsoft Powerpoint (microsoft.com), and Adobe Photoshop (adobe.com). Plots were produced using Matplotlib.^[60]^

## Acknowledgements

The authors thank all members of the Gonen lab for helpful and critical discussions. This research was supported by the Department of Defense HDTRA1-21-1-0004 and The National Institute of General Medical Sciences, grant R35-GM142797.

