## Supplementary Figures and Table for "δ-Conotoxin Structure Prediction and Analysis through Large-scale Comparative and Deep Learning Modeling Approaches"

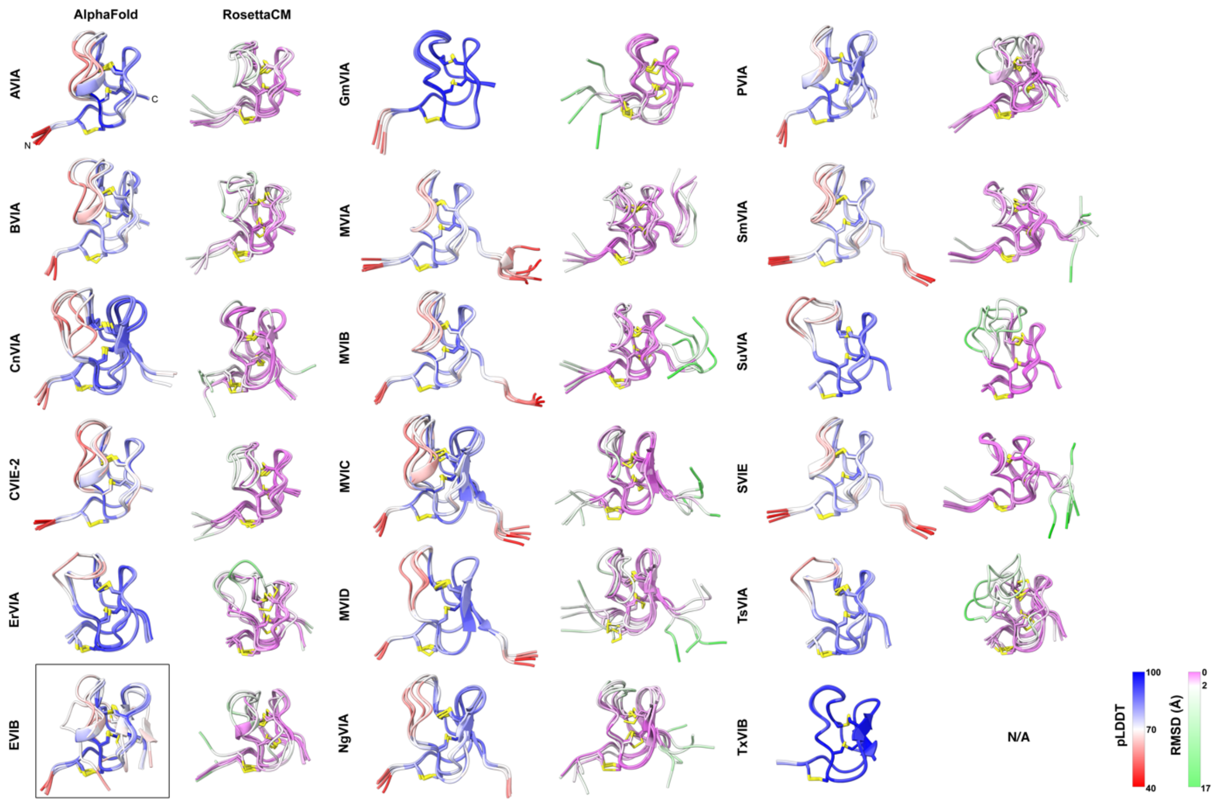


**Figure S1.** Results of the modeling pipeline for all peptides. From left: all five models produced by AlphaFold, colored by per-residue pLDDT score; representatives of the top 5 best-scoring clusters produced by RosettaCM, colored by per-residue RMSD. The sole model generated which did not meet the disulfide bonding criterion is visible in the black box together with the four other models having the correct ICK fold. RMSD and pLDDT color bars are shown bottom right.

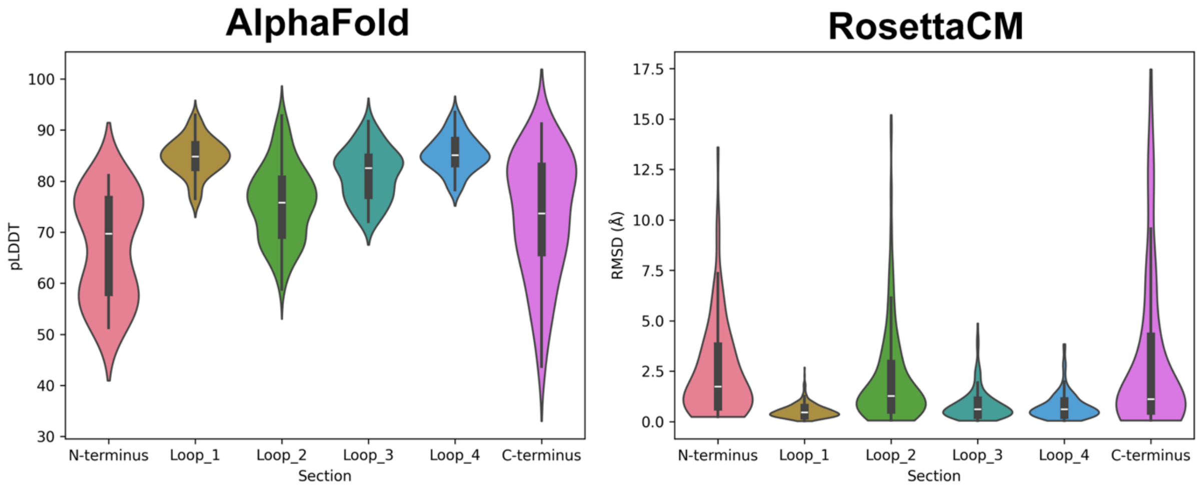


**Figure S2.** Violin plots indicating (left) mean per-residue pLDDT scores and (right) mean per-residue RMSD for different regions of the peptide models in this study. The termini and loop 2 regions show higher RMSD and lower pLDDT scores than loops 1, 3 and 4.

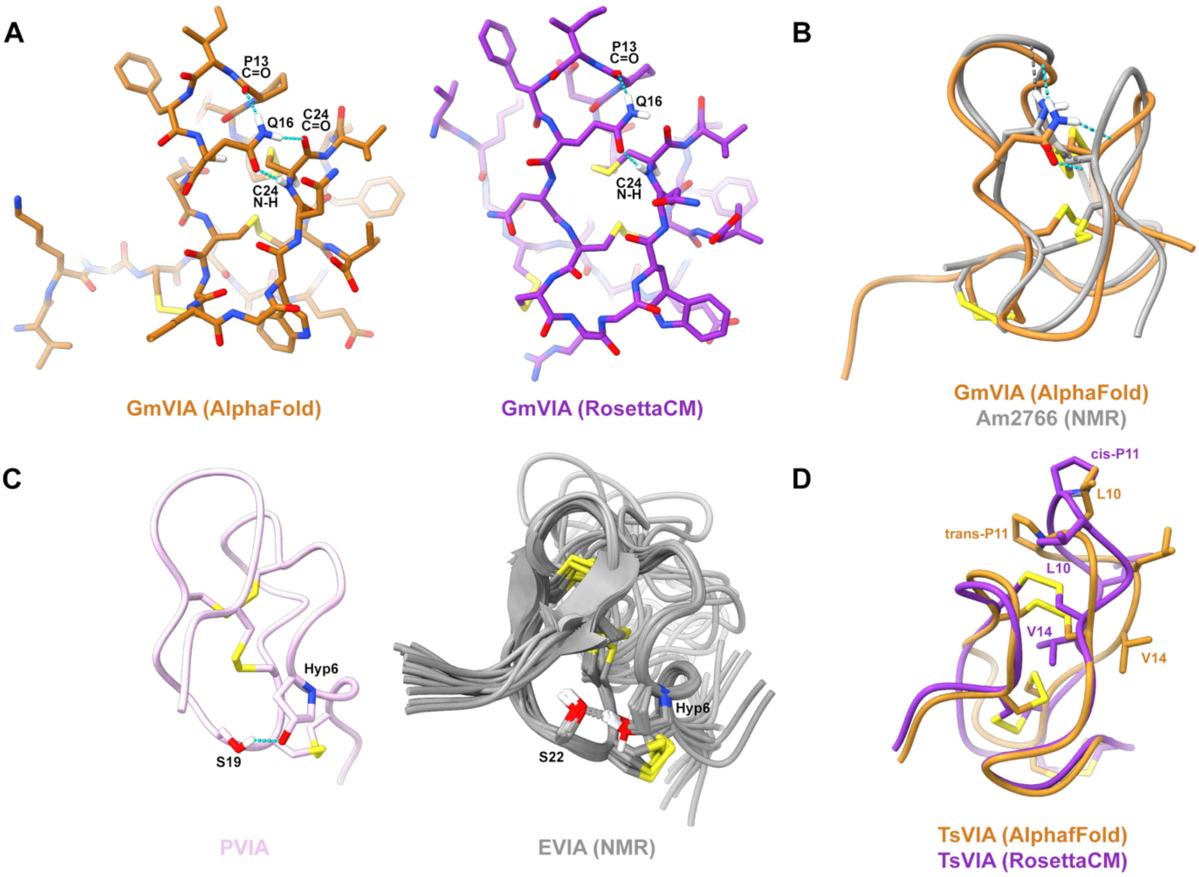


**Figure S3**. A) Top-scoring AlphaFold (left) and RosettaCM (right) models of GmVIA highlighting the hydrogen bonding of the Q16 sidechain with backbone atoms on adjacent loops B) Comparison with a similar hydrogen-bonding pattern in one NMR-derived structure of Am2766 C) RosettaCM model of PVIA (left) showing the hydrogen bonding interaction between the sidechains of S19 and a conserved Hyp residue, and (right) a similar bonding pattern observed in the experimental ensemble of structures of EVIA derived by NMR. D) Comparison of the top-scoring AlphaFold (orange) and RosettaCM (magenta) models of TsVIA, showing the effect of the cis-proline bond on the position of nearby sidechains on loop 2.

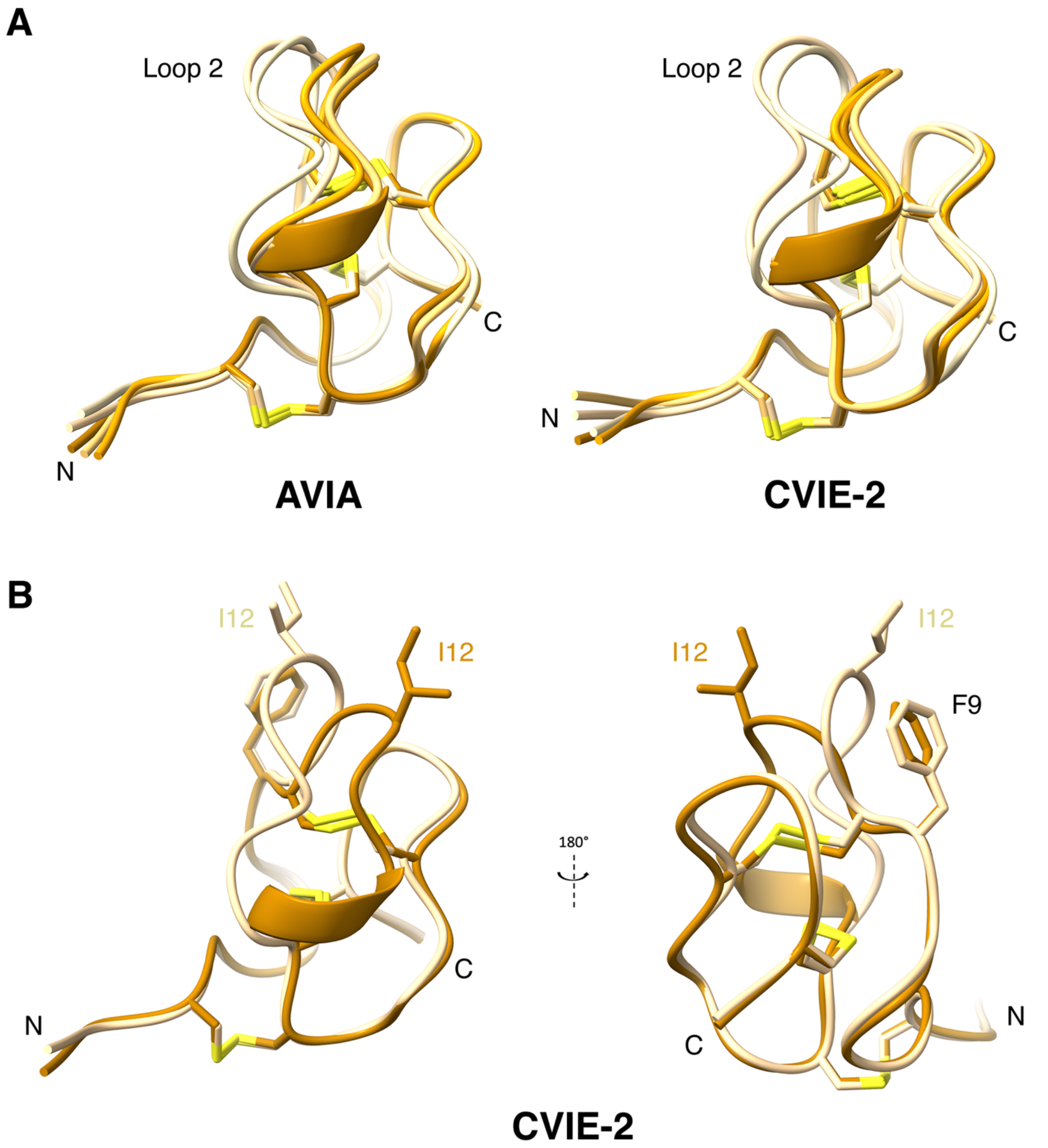


**Figure S4**. Selected features of the δ-conotoxin models. A) Top 5 cluster representatives from AlphaFold modeling of (left) AVIA and (right) CVIE-2, bifurcation of loop 2. Models are colored from best-scoring (dark orange) to worst (white). Disulfide bonds are highlighted in yellow. B) Two views of two AlphaFold models of CVIE-2 illustrating the positioning of the sidechains of putative bioactive residues I12 and F9 in the two main positions of loop 2.

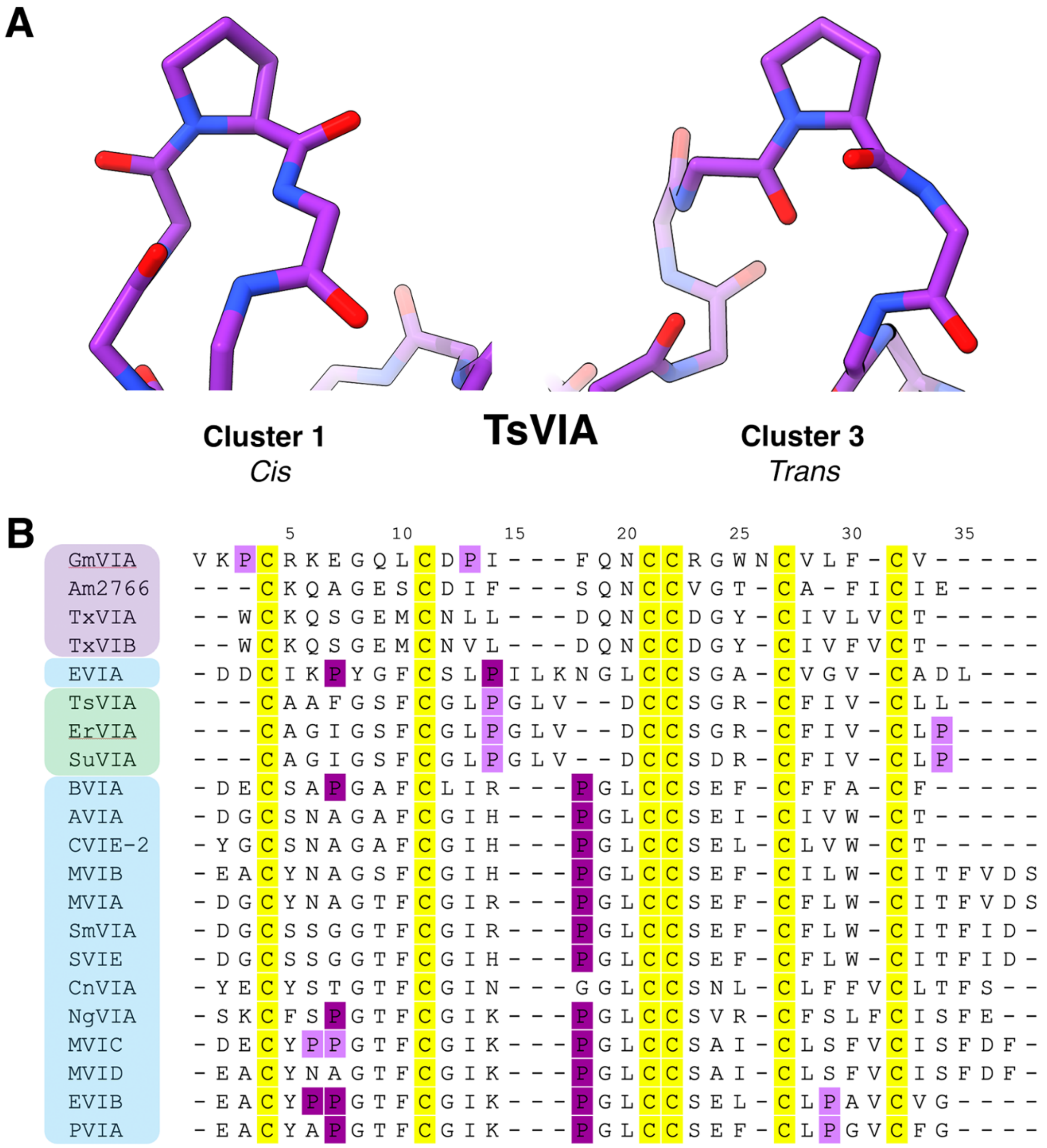


**Figure S5**. A) Comparison of representatives of two clusters from the RosettaCM modeling of TsVIA, showing the presence of a cis proline in cluster 1 but a trans proline at the same position in cluster 3. B) Sequence alignment of the peptides in this study, with proline residues highlighted in purple. Prolines modeled as hydroxylated in the RosettaCM pipeline are indicated in dark purple.

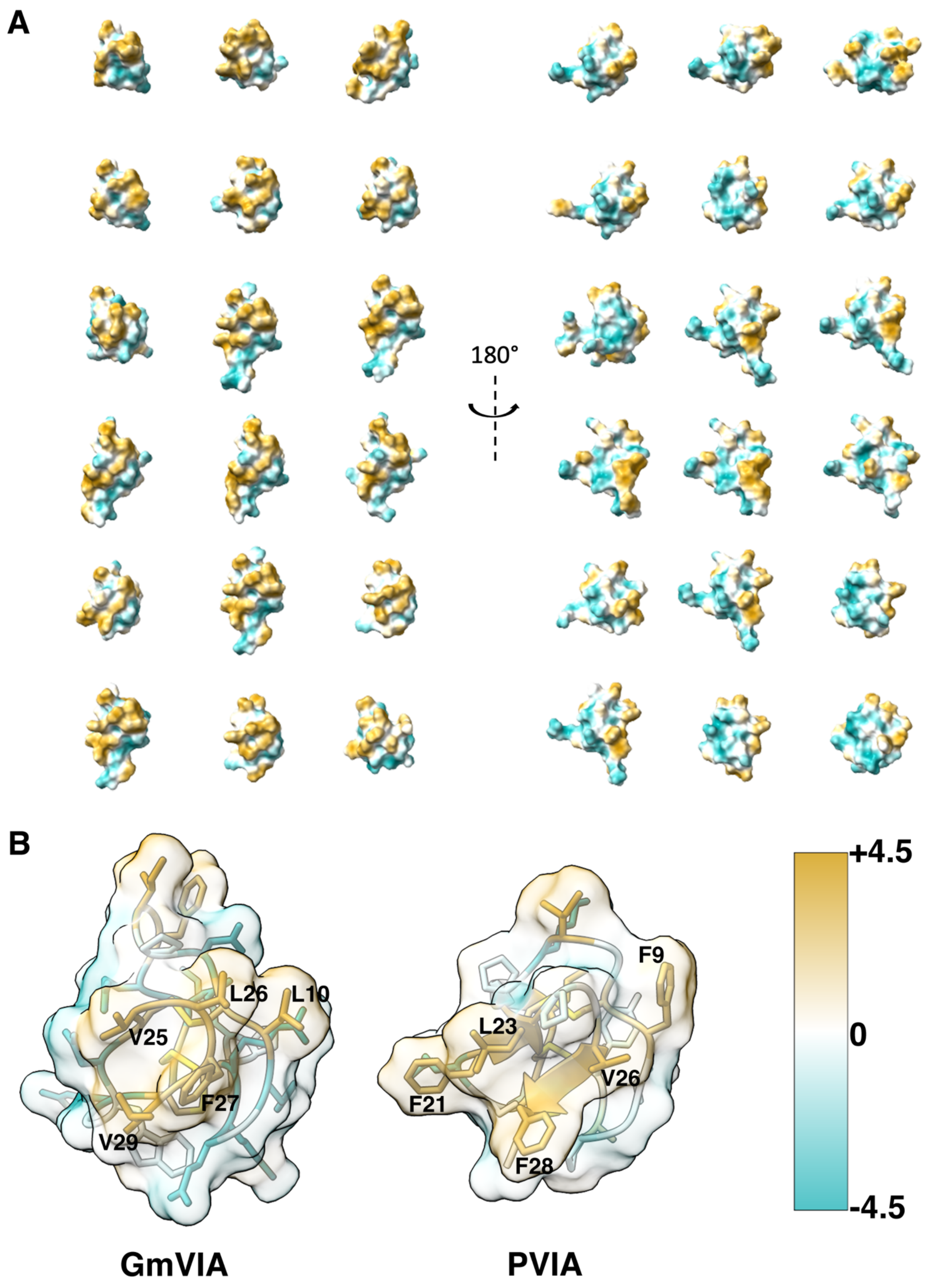


**Figure S6**. A) Surfaces of the top-scoring AlphaFold models for the peptides in this study (displayed alphabetically from top left) colored by Kyte-Doolittle hydrophobicity,[61] showing (left) the consensus hydrophobic patch and (right) the reverse side. B) Enlarged views of

GmVIA and PVIA highlighting the key residues responsible for forming the hydrophobic patch. Hydrophobicity scale color bar is shown bottom right.

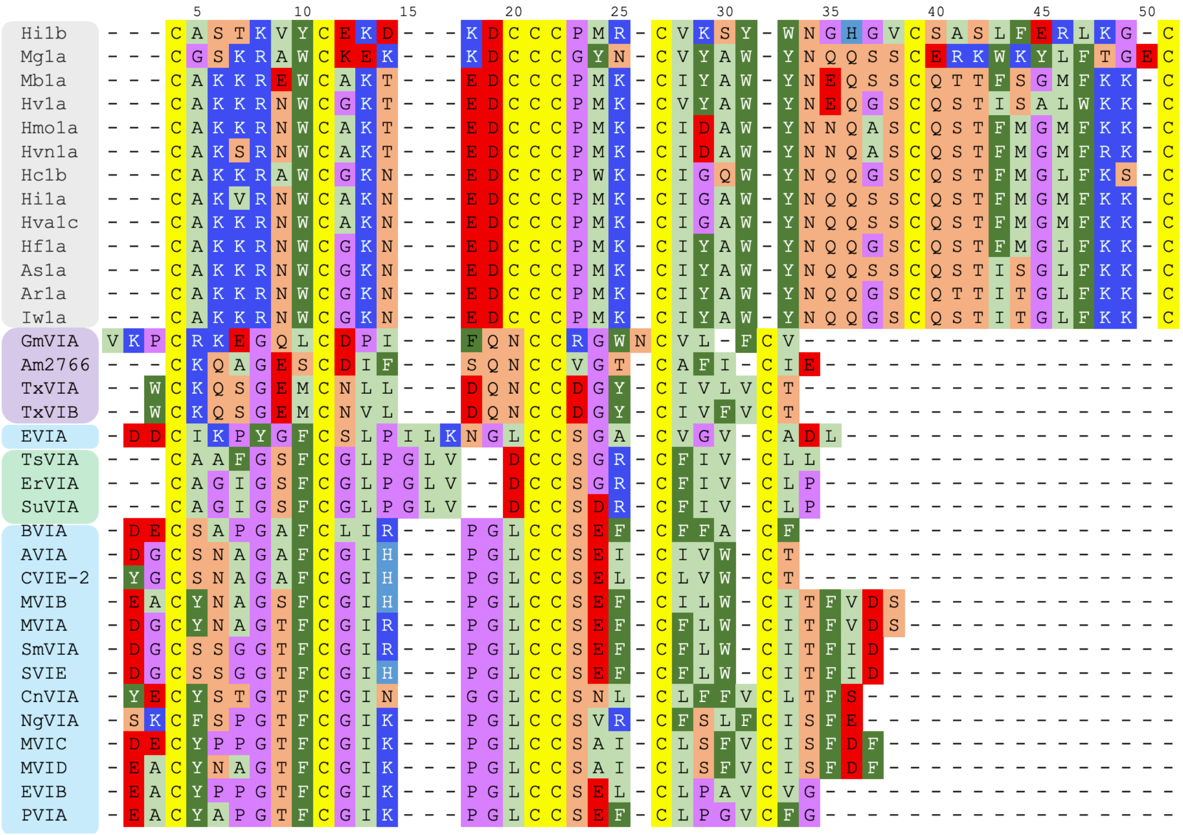


**Figure S7**. Sequence alignment of all the δ-conotoxins in this study (color scheme as for Figure 1) with 14 representative δ-hexatoxin sequences (grey) determined by Herzig et al.[28]

| NMR Structure | Clashscore | Poor rotamers | Favored rotamers | Ramachandran outliers | Ramachandran favored | Rama distribution Z-score | Molprobity score | Cis-Pro | Cis-nonPro | Twisted peptides |
|---|---|---|---|---|---|---|---|---|---|---|
| 1fu3_model_1 | 83.54 | 8 | 10 | 1 | 17 | -5.17 ± 1.19 | 4.39 | 0 | 0 | 0 |
| 1fu3_model_2 | 96.20 | 7 | 12 | 2 | 15 | -7.45 ± 0.74 | 4.46 | 0 | 0 | 0 |
| 1fu3_model_3 | 91.14 | 11 | 7 | 3 | 13 | -7.03 ± 1.01 | 4.64 | 0 | 0 | 0 |
| 1fu3_model_4 | 91.14 | 9 | 9 | 3 | 15 | -5.40 ± 1.33 | 4.52 | 0 | 0 | 0 |
| 1fu3_model_5 | 101.27 | 11 | 7 | 1 | 15 | -4.60 ± 1.78 | 4.63 | 0 | 0 | 0 |
| 1fu3_model_6 | 108.86 | 13 | 9 | 1 | 16 | -7.26 ± 0.65 | 4.69 | 0 | 0 | 0 |
| 1fu3_model_7 | 78.48 | 8 | 10 | 3 | 17 | -5.89 ± 1.12 | 4.36 | 0 | 0 | 0 |
| 1fu3_model_8 | 83.54 | 10 | 9 | 1 | 16 | -7.32 ± 0.72 | 4.49 | 0 | 0 | 0 |
| 1fu3_model_9 | 81.01 | 8 | 11 | 3 | 16 | -5.26 ± 1.26 | 4.41 | 0 | 0 | 0 |
| 1fu3_model_10 | 73.42 | 9 | 11 | 1 | 14 | -7.22 ± 0.77 | 4.46 | 0 | 0 | 0 |
| 1fu3_model_11 | 93.67 | 9 | 11 | 2 | 14 | -6.89 ± 0.93 | 4.56 | 0 | 0 | 0 |
| 1fu3_model_12 | 101.27 | 6 | 13 | 2 | 18 | -6.14 ± 0.97 | 4.34 | 0 | 0 | 0 |
| 1fu3_model_13 | 96.20 | 11 | 8 | 1 | 18 | -6.25 ± 1.05 | 4.52 | 0 | 0 | 0 |
| 1fu3_model_14 | 81.01 | 8 | 11 | 1 | 17 | -5.71 ± 1.16 | 4.38 | 0 | 0 | 0 |
| 1fu3_model_15 | 88.61 | 10 | 10 | 1 | 18 | -5.64 ± 1.14 | 4.45 | 0 | 0 | 0 |
| 1fu3_model_16 | 83.54 | 7 | 13 | 2 | 16 | -5.27 ± 1.41 | 4.38 | 0 | 0 | 0 |
| 1fu3_model_17 | 96.20 | 5 | 13 | 2 | 16 | -6.80 ± 1.00 | 4.33 | 0 | 0 | 0 |
| 1fu3_model_18 | 88.61 | 8 | 11 | 3 | 14 | -7.14 ± 0.91 | 4.50 | 0 | 0 | 0 |
| 1fu3_model_19 | 75.95 | 9 | 9 | 3 | 17 | -7.28 ± 0.79 | 4.39 | 0 | 0 | 0 |
| 1fu3_model_20 | 118.99 | 8 | 12 | 1 | 16 | -5.87 ± 1.08 | 4.57 | 0 | 0 | 0 |
| 1yz2_model_1 | 64.25 | 8 | 7 | 0 | 18 | -6.31 ± 0.85 | 4.26 | 0 | 0 | 0 |
| 1yz2_model_2 | 47.49 | 8 | 8 | 1 | 18 | -3.67 ± 1.62 | 4.13 | 0 | 0 | 0 |
| 1yz2_model_3 | 47.49 | 7 | 7 | 1 | 18 | -6.12 ± 1.10 | 4.09 | 0 | 0 | 0 |
| 1yz2_model_4 | 55.87 | 11 | 6 | 0 | 18 | -5.47 ± 1.13 | 4.30 | 0 | 0 | 0 |
| 1yz2_model_5 | 44.69 | 9 | 4 | 0 | 15 | -6.30 ± 1.18 | 4.25 | 0 | 0 | 0 |
| 1yz2_model_6 | 55.87 | 6 | 6 | 1 | 15 | -6.81 ± 0.96 | 4.21 | 0 | 0 | 0 |

| | | | | | | | | | | |
|---|---|---|---|---|---|---|---|---|---|---|
| 1yz2_model_7 | 64.25 | 8 | 4 | 1 | 18 | -5.48 ± 1.26 | 4.26 | 0 | 0 | 0 |
| 1yz2_model_8 | 55.87 | 10 | 8 | 1 | 19 | -4.68 ± 1.44 | 4.23 | 0 | 0 | 0 |
| 1yz2_model_9 | 55.87 | 9 | 9 | 1 | 16 | -5.50 ± 1.25 | 4.31 | 0 | 0 | 0 |
| 1yz2_model_10 | 58.66 | 5 | 11 | 0 | 17 | -6.43 ± 1.12 | 4.11 | 0 | 0 | 0 |
| 1yz2_model_11 | 36.31 | 9 | 10 | 0 | 15 | -5.83 ± 1.47 | 4.16 | 0 | 0 | 0 |
| 1yz2_model_12 | 41.90 | 9 | 7 | 0 | 17 | -5.24 ± 1.52 | 4.16 | 0 | 0 | 0 |
| 1yz2_model_13 | 53.07 | 9 | 6 | 2 | 17 | -5.12 ± 1.34 | 4.26 | 0 | 0 | 0 |
| 1yz2_model_14 | 50.28 | 7 | 7 | 0 | 18 | -6.30 ± 1.09 | 4.11 | 0 | 0 | 0 |
| 1yz2_model_15 | 55.87 | 8 | 8 | 1 | 18 | -5.44 ± 1.14 | 4.20 | 0 | 0 | 0 |
| 1g1p_model_1 | 0.00 | 1 | 22 | 0 | 21 | -4.91 ± 0.93 | 1.72 | 0 | 0 | 0 |
| 1g1p_model_2 | 2.26 | 1 | 20 | 1 | 19 | -5.20 ± 1.01 | 2.30 | 0 | 0 | 0 |
| 1g1p_model_3 | 0.00 | 1 | 22 | 1 | 19 | -5.35 ± 0.86 | 1.80 | 0 | 0 | 0 |
| 1g1p_model_4 | 2.26 | 1 | 20 | 1 | 20 | -5.65 ± 0.76 | 2.26 | 0 | 0 | 0 |
| 1g1p_model_5 | 2.26 | 0 | 18 | 1 | 23 | -5.27 ± 0.85 | 1.66 | 0 | 0 | 0 |
| 1g1p_model_6 | 0.00 | 1 | 20 | 1 | 22 | -5.99 ± 0.63 | 1.67 | 0 | 0 | 0 |
| 1g1p_model_7 | 0.00 | 1 | 17 | 2 | 16 | -6.12 ± 0.92 | 1.88 | 0 | 0 | 0 |
| 1g1p_model_8 | 0.00 | 1 | 20 | 0 | 19 | -6.38 ± 0.80 | 1.80 | 0 | 0 | 0 |
| 1g1p_model_9 | 0.00 | 2 | 18 | 3 | 20 | -5.39 ± 1.00 | 1.99 | 0 | 0 | 0 |
| 1g1p_model_10 | 0.00 | 1 | 17 | 3 | 19 | -6.01 ± 1.09 | 1.80 | 0 | 0 | 0 |
| 1g1p_model_11 | 0.00 | 0 | 22 | 2 | 19 | -6.29 ± 0.81 | 1.34 | 0 | 0 | 0 |
| 1g1p_model_12 | 2.26 | 1 | 17 | 2 | 17 | -6.20 ± 1.02 | 2.36 | 0 | 0 | 0 |
| 1g1p_model_13 | 0.00 | 0 | 19 | 1 | 19 | -5.89 ± 0.89 | 1.34 | 0 | 0 | 0 |
| 1g1p_model_14 | 0.00 | 2 | 22 | 1 | 21 | -4.66 ± 1.08 | 1.95 | 0 | 0 | 0 |
| 1g1p_model_15 | 0.00 | 2 | 20 | 1 | 22 | -5.27 ± 0.87 | 1.90 | 0 | 0 | 0 |
| 1g1p_model_16 | 0.00 | 0 | 18 | 2 | 17 | -5.20 ± 0.92 | 1.40 | 0 | 0 | 0 |
| 1g1p_model_17 | 0.00 | 2 | 19 | 1 | 20 | -4.72 ± 1.10 | 1.99 | 0 | 0 | 0 |
| 1g1p_model_18 | 0.00 | 0 | 20 | 2 | 21 | -4.68 ± 1.12 | 1.26 | 0 | 0 | 0 |

| Model | Clashscore | Poor rotamers | Favored rotamers | Ramachandran outliers | Ramachandran favored | Rama distribution Z-score | Molprobity score | Cis-Pro | Cis-nonPro | Twisted peptides |
|---|---|---|---|---|---|---|---|---|---|---|
| AVIA_c.1.1 | 0.00 | 0 | 20 | 0 | 20 | 1.29 ± 1.83 | 1.02 | 0 | 1 | 0 |
| AVIA_beta | 0.00 | 0 | 19 | 0 | 18 | 0.13 ± 1.91 | 1.21 | 0 | 1 | 0 |
| AVIA_ref2015 | 2.81 | 0 | 20 | 0 | 19 | -2.64 ± 1.41 | 1.70 | 0 | 1 | 0 |
| AVIA_ranked0 | 5.63 | 2 | 18 | 2 | 22 | -1.65 ± 1.76 | 2.65 | 0 | 0 | 0 |
| BVIA_c.1.1 | 0.00 | 0 | 20 | 0 | 17 | -4.07 ± 1.44 | 1.06 | 0 | 0 | 0 |
| BVIA_beta | 0.00 | 0 | 20 | 1 | 16 | -3.96 ± 1.44 | 1.17 | 0 | 0 | 0 |
| BVIA_ref2015 | 0.00 | 0 | 20 | 0 | 17 | -3.53 ± 1.35 | 1.06 | 0 | 0 | 0 |
| BVIA_ranked0 | 2.63 | 1 | 18 | 0 | 21 | -1.58 ± 1.44 | 2.22 | 0 | 0 | 0 |
| CnVIA_c.1.1 | 0.00 | 0 | 26 | 0 | 27 | -0.71 ± 1.29 | 0.74 | 0 | 0 | 0 |
| CnVIA_beta | 2.32 | 0 | 26 | 0 | 28 | -1.11 ± 1.18 | 1.01 | 0 | 1 | 0 |
| CnVIA_ref2015 | 0.00 | 0 | 26 | 0 | 27 | -1.44 ± 1.35 | 0.74 | 0 | 1 | 0 |
| CnVIA_ranked0 | 2.32 | 1 | 22 | 0 | 27 | -2.53 ± 1.21 | 1.69 | 0 | 0 | 0 |
| CVIE-2.c.1.1 | 0.00 | 0 | 20 | 0 | 19 | 1.24 ± 2.19 | 1.13 | 0 | 1 | 0 |
| CVIE-2_beta | 2.74 | 0 | 20 | 1 | 19 | 0.72 ± 2.03 | 1.70 | 0 | 1 | 0 |
| CVIE-2_ref2015 | 0.00 | 0 | 20 | 0 | 18 | -2.75 ± 1.74 | 1.21 | 0 | 1 | 0 |
| CVIE-2_ranked0 | 13.74 | 6 | 13 | 2 | 23 | -5.23 ± 0.70 | 3.24 | 0 | 0 | 0 |
| ErVIA_c.1.1 | 0.00 | 0 | 21 | 1 | 22 | -0.60 ± 1.72 | 1.10 | 0 | 0 | 0 |
| ErVIA_beta | 0.00 | 0 | 21 | 0 | 22 | 2.26 ± 2.11 | 1.10 | 0 | 0 | 0 |
| ErVIA_ref2015 | 0.00 | 0 | 21 | 0 | 22 | -2.63 ± 1.39 | 1.10 | 0 | 0 | 0 |
| ErVIA_ranked0 | 0.00 | 1 | 17 | 1 | 24 | -3.22 ± 1.07 | 1.29 | 0 | 0 | 0 |
| EVIB_c.1.1 | 2.51 | 0 | 20 | 0 | 17 | -2.11 ± 1.63 | 1.69 | 0 | 1 | 0 |
| EVIB_beta | 0.00 | 0 | 20 | 0 | 18 | -3.14 ± 1.28 | 1.05 | 0 | 1 | 0 |
| EVIB_ref2015 | 5.03 | 0 | 20 | 0 | 19 | -1.25 ± 1.80 | 1.61 | 0 | 0 | 0 |
| EVIB_ranked0 | 0.00 | 1 | 22 | 3 | 24 | -3.27 ± 1.39 | 1.56 | 0 | 0 | 1 |
| GmVIA_c.1.1 | 2.20 | 0 | 27 | 0 | 25 | -1.70 ± 1.34 | 1.46 | 0 | 0 | 0 |
| GmVIA_beta | 0.00 | 0 | 27 | 0 | 25 | -0.37 ± 1.47 | 0.96 | 0 | 0 | 0 |

| | | | | | | | | | | |
|---|---|---|---|---|---|---|---|---|---|---|
| GmVIA_ref2015 | 0.00 | 0 | 27 | 0 | 25 | -0.80 ± 1.25 | 0.96 | 0 | 0 | 0 |
| GmVIA_ranked0 | 6.59 | 1 | 26 | 1 | 25 | -0.36 ± 1.41 | 2.26 | 0 | 0 | 0 |
| MVIA_c.1.1 | 6.49 | 0 | 26 | 0 | 24 | 1.41 ± 1.87 | 1.94 | 0 | 1 | 0 |
| MVIA_beta | 2.16 | 0 | 26 | 0 | 24 | -0.58 ± 1.77 | 1.57 | 0 | 1 | 0 |
| MVIA_ref2015 | 4.33 | 0 | 26 | 0 | 24 | -1.21 ± 1.51 | 1.79 | 0 | 0 | 0 |
| MVIA_ranked0 | 4.36 | 4 | 21 | 2 | 26 | -3.49 ± 1.46 | 2.73 | 0 | 0 | 1 |
| MVIB_c.1.1 | 4.38 | 0 | 26 | 0 | 23 | 1.84 ± 1.89 | 1.87 | 0 | 1 | 0 |
| MVIB_beta | 2.19 | 0 | 26 | 1 | 25 | 2.27 ± 1.77 | 1.46 | 0 | 1 | 0 |
| MVIB_ref2015 | 4.38 | 0 | 26 | 0 | 25 | -1.77 ± 1.39 | 1.68 | 0 | 1 | 0 |
| MVIB_ranked0 | 4.41 | 2 | 23 | 2 | 26 | -2.95 ± 1.48 | 2.51 | 0 | 0 | 0 |
| MVIC_c.1.1 | 0.00 | 0 | 27 | 0 | 24 | -1.65 ± 1.42 | 1.08 | 0 | 1 | 0 |
| MVIC_beta | 0.00 | 0 | 27 | 0 | 26 | -0.37 ± 1.46 | 0.75 | 0 | 1 | 0 |
| MVIC_ref2015 | 2.19 | 0 | 27 | 0 | 23 | -1.46 ± 1.43 | 1.65 | 0 | 1 | 0 |
| MVIC_ranked0 | 0.00 | 1 | 25 | 0 | 27 | -2.45 ± 1.33 | 1.47 | 0 | 0 | 0 |
| MVID_c.1.1 | 0.00 | 0 | 25 | 0 | 24 | 0.12 ± 1.69 | 1.08 | 0 | 1 | 0 |
| MVID_beta | 0.00 | 0 | 25 | 0 | 27 | 0.11 ± 1.48 | 0.50 | 0 | 0 | 0 |
| MVID_ref2015 | 0.00 | 0 | 25 | 0 | 25 | -2.02 ± 1.34 | 0.96 | 0 | 0 | 0 |
| MVID_ranked0 | 0.00 | 0 | 25 | 0 | 27 | -3.97 ± 1.06 | 1.05 | 0 | 0 | 0 |
| NgVIA_c.1.1 | 0.00 | 0 | 26 | 0 | 21 | -0.17 ± 1.43 | 1.01 | 0 | 1 | 0 |
| NgVIA_beta | 0.00 | 0 | 26 | 0 | 21 | -0.94 ± 1.54 | 1.01 | 0 | 1 | 0 |
| NgVIA_ref2015 | 4.37 | 0 | 26 | 0 | 20 | -0.61 ± 1.38 | 1.84 | 0 | 0 | 0 |
| NgVIA_ranked0 | 2.19 | 1 | 25 | 2 | 24 | -2.73 ± 1.41 | 2.11 | 0 | 0 | 0 |
| PVIA_c.1.1 | 0.00 | 0 | 20 | 0 | 18 | -1.72 ± 1.98 | 1.15 | 0 | 1 | 0 |
| PVIA_beta | 2.53 | 0 | 20 | 0 | 19 | -0.09 ± 1.91 | 1.57 | 0 | 0 | 0 |
| PVIA_ref2015 | 10.13 | 0 | 20 | 0 | 18 | -4.87 ± 0.99 | 2.17 | 0 | 0 | 0 |
| PVIA_ranked0 | 0.00 | 3 | 18 | 2 | 24 | -2.30 ± 1.50 | 1.94 | 0 | 0 | 1 |
| SmVIA_c.1.1 | 0.00 | 0 | 25 | 0 | 23 | -1.86 ± 1.93 | 1.09 | 0 | 1 | 0 |
| SmVIA_beta | 0.00 | 0 | 25 | 0 | 23 | -1.51 ± 1.49 | 1.09 | 0 | 2 | 0 |

| | | | | | | | | | | |
|---|---|---|---|---|---|---|---|---|---|---|
| SmVIA_ref2015 | 4.59 | 0 | 25 | 0 | 23 | -1.47 ± 1.95 | 1.82 | 0 | 1 | 0 |
| SmVIA_ranked0 | 4.60 | 3 | 20 | 2 | 24 | -5.21 ± 0.96 | 2.74 | 0 | 0 | 1 |
| SuVIA_c.1.1 | 2.73 | 0 | 22 | 0 | 24 | 0.62 ± 1.78 | 1.33 | 0 | 0 | 0 |
| SuVIA_beta | 0.00 | 0 | 22 | 0 | 23 | 1.24 ± 1.66 | 0.99 | 0 | 0 | 0 |
| SuVIA_ref2015 | 5.46 | 0 | 22 | 0 | 21 | 0.49 ± 1.73 | 1.97 | 0 | 0 | 0 |
| SuVIA_ranked0 | 0.00 | 1 | 19 | 1 | 21 | -2.89 ± 1.55 | 1.68 | 0 | 0 | 0 |
| SVIE_c.1.1 | 0.00 | 0 | 25 | 0 | 24 | -0.76 ± 1.80 | 0.97 | 0 | 1 | 0 |
| SVIE_beta | 2.33 | 0 | 25 | 0 | 23 | -2.17 ± 1.39 | 1.60 | 0 | 1 | 0 |
| SVIE_ref2015 | 4.66 | 0 | 25 | 0 | 22 | -0.99 ± 1.97 | 1.90 | 0 | 1 | 0 |
| SVIE_ranked0 | 4.67 | 3 | 20 | 2 | 25 | -4.51 ± 1.08 | 2.68 | 0 | 0 | 0 |
| TsVIA_c.1.1 | 0.00 | 0 | 21 | 1 | 21 | -2.24 ± 1.43 | 1.18 | 1 | 0 | 0 |
| TsVIA_beta | 2.70 | 0 | 21 | 0 | 23 | 2.45 ± 1.92 | 1.54 | 0 | 0 | 0 |
| TsVIA_ref2105 | 0.00 | 0 | 21 | 0 | 21 | -0.73 ± 1.62 | 1.18 | 0 | 0 | 0 |
| TsVIA_ranked0 | 5.41 | 3 | 14 | 1 | 22 | -1.81 ± 1.60 | 2.77 | 0 | 0 | 0 |
| 1fu3_mut_0035 | 2.54 | 0 | 25 | 1 | 22 | 0.05 ± 1.65 | 1.64 | 0 | 0 | 0 |
| TxVIB_ranked0 | 7.63 | 1 | 22 | 0 | 22 | -0.62 ± 1.46 | 2.47 | 0 | 0 | 0 |

**Table S1**. Selected geometrical statistics for NMR ensemble structures and models discussed in this work, as calculated by MolProbity. Model or structure names are colored according to the source cone snail: blue (piscivorous), green (vermivorous) and magenta (molluscivorous). The top-scoring RosettaCM model is designated ####_c.1.1; the top-scoring AlphaFold model is designated ####_ranked0. The top-scoring models produced by the RosettaCM workflow using the ref2015_cart and beta_nov16_cart scorefunctions are designated ####_ref2015 and ####_beta, respectively.
